# An elastic proteinaceous envelope encapsulates the early *Arabidopsis* embryo

**DOI:** 10.1101/2023.05.03.539186

**Authors:** Yosapol Harnvanichvech, Cecilia Borassi, Diaa Eldin S. Daghma, Hanne M. van der Kooij, Joris Sprakel, Dolf Weijers

## Abstract

Plant external surfaces are often covered by barriers that control the exchange of molecules, protect from pathogens, and offer mechanical integrity. A key question is when and how such surface barriers are generated. Post-embryonic surfaces have well-studied barriers, including the cuticle, and late Arabidopsis embryo was shown to be protected by an endosperm-derived sheath deposited onto a primordial cuticle. Here we show that both cuticle and sheath are preceded by another structure during the earliest stages of embryogenesis. This structure, which we named the embryonic envelope, is tightly wrapped around the embryonic surface but can be physically detached by cell wall digestion. We show that this structure is composed primarily of Extensin and Arabinogalactan *O*-glycoproteins and lipids, which appear to form a dense and elastic crosslinked embryonic envelope. The envelope forms in cuticle-deficient mutants and in a mutant that lacks endosperm. This embryo-derived envelope is therefore distinct from previously described cuticle and sheath structures. We propose that it acts as an expandable diffusion barrier, as well as a means to mechanically confine the embryo to maintain its tensegrity during early embryogenesis.

**Summary statement:** The early Arabidopsis embryo is surrounded by a proteinaceous envelope that is distinct from the cuticle or embryo sheath

## Introduction

Cells in land plants are generally exposed to the atmosphere, and are often covered by structures that prevent evaporation, exchange of solutes, and invasion of pathogens (Nawrath et al., 2013; Yeats et al., 2013). These structures include the cuticle (Nawrath 2006, Nawrath et al., 2013), as well as suberin lamellae (Nawrath et al., 2013). In seed plants, the seedling develops from an embryo that is covered by seed coat tissues during its development (Ingouff et al., 2006; Voiniciuc et al., 2015; Sechet et al., 2018), and the cuticle needs to be complete before seed germination (De Giorgi et al., 2015; Coen et al, 2019; Creff et al., 2019). In flowering plants, the embryo shares the seed cavity with the product of a second, independent fertilization event, the endosperm (Berger et al., 2003; Hamamura et al., 2012; Ali et al., 2023). As such, the flowering plant embryo surface is exposed to an external environment well before germination, and it is not yet clear to what extent the exchange of materials at the epidermal interface is controlled by a barrier during embryo development. Also, the absence of direct mechanical constraints imposed by the seed coat may urge the need for another structure to confine the embryo during its development. In recent years, a structure has been described to cover the Arabidopsis embryo surface (Fourquin et al., 2016; Moussu et al., 2017; Doll & Ingram, 2020). After double fertilization the development of the endosperm and the embryo starts, and optimal seed development and germination depend on the communication at the interface between these tissues. At later stages of seed development, the endosperm cellularizes, while the embryo grows invasively into it. At this point, a selective apoplastic lipid-rich barrier, the embryo cuticle, is secreted by the embryonic epidermal cells and limits the communication between the embryo and the endosperm. At late globular stage the secretion of cuticle components in vesicles (cutinosomes) starts, arranging in patches that begin to assemble on the embryo proper surface, and by heart stage it is observed as a continuous layer (Creff et al., 2019). Some molecular components have been identified as key players for the proper assembly of the cutinosomes in a mechanism that requires signaling between the endosperm and the embryo. This pathway involves the endosperm-specific subtilisin protease Abnormal Leaf-shape 1 (ALE1) and the embryo-expressed receptor kinases GASSHO1 (GSO1) and GASSHO2 (GSO2) (Creff et al., 2019). Later, the embryo continues to grow and the endosperm is progressively degraded. From the heart stage onwards, endosperm-derived materials accumulate to form a structure on the outer face of the embryo cuticle, rich in Extensin proteins, known as the embryo sheath. This structure is required for normal embryo development preventing its adhesion to the endosperm during invasive growth. It has been shown that the sheath enables the cotyledons to break through the seed coat and it acts as the first barrier between the aerial organs and the environment (Doll et al., 2020).

An open question is whether any structure precedes the formation of the embryonic cuticle and embryonic sheath. It could be that no such structure is needed at early stages, but it might well be that also at the earliest stages, an extracellular structure controls exchange between embryo and endosperm. Also, since turgor pressure in endosperm is dynamic throughout seed development (Creff et al., 2023) the embryo may need an extracellular structure to keep its tensegrity.

We describe the existence of a previously undescribed extracellular envelope that exists on the surface of the Arabidopsis embryo well before the formation of cuticle and sheath. The envelope is a protein- and lipid-rich, embryo-derived structure and its biogenesis depends on the determination of epidermal cell fate. We speculate that the formation of the embryonic envelope could serve as a diffusion barrier that both chemically and physically separates the early embryo from the developing endosperm prior to the assembly of the embryonic cuticle.

## Results

### Early *Arabidopsis* embryos are surrounded by an extracellular structure

We hypothesized that a structure with barrier-like properties should surround the Arabidopsis embryo at early stages. This structure would envelope the embryo, acting as a separation layer between the embryo and the developing endosperm and it should be flexible enough to accommodate rapid growth. We first performed transmission electron microscopy (TEM) on wild-type seeds. To enable high-definition analysis of native structures, we optimized a method based on encapsulating individual seeds in a mixture of yeast and cyanobacteria (filler), followed by high-pressure freezing and freeze-substitution (Daghma et al., in preparation). This method allows ultra-rapid immobilization of cellular processes, thus enabling the study of 3D organization of subcellular structures. The excellent structural preservation and high resolution allowed a detailed analysis of the embryo surface. Importantly, no rupture of the endosperm-embryo interface occurred during processing. At both 8-cell and globular stages, we detected a continuous layer surrounding both proembryo and suspensor that appeared electron-dense. The layer appeared homogeneous along the embryo surface. We measured cross-sectional thickness in sections from 5 different embryos. The thickness around the proembryo and along the suspensor ranged between 40-70 nm (Figure 1). Hence the structure is thicker than the plasma membrane but thinner than a primary cell wall.

**Figure 1.**
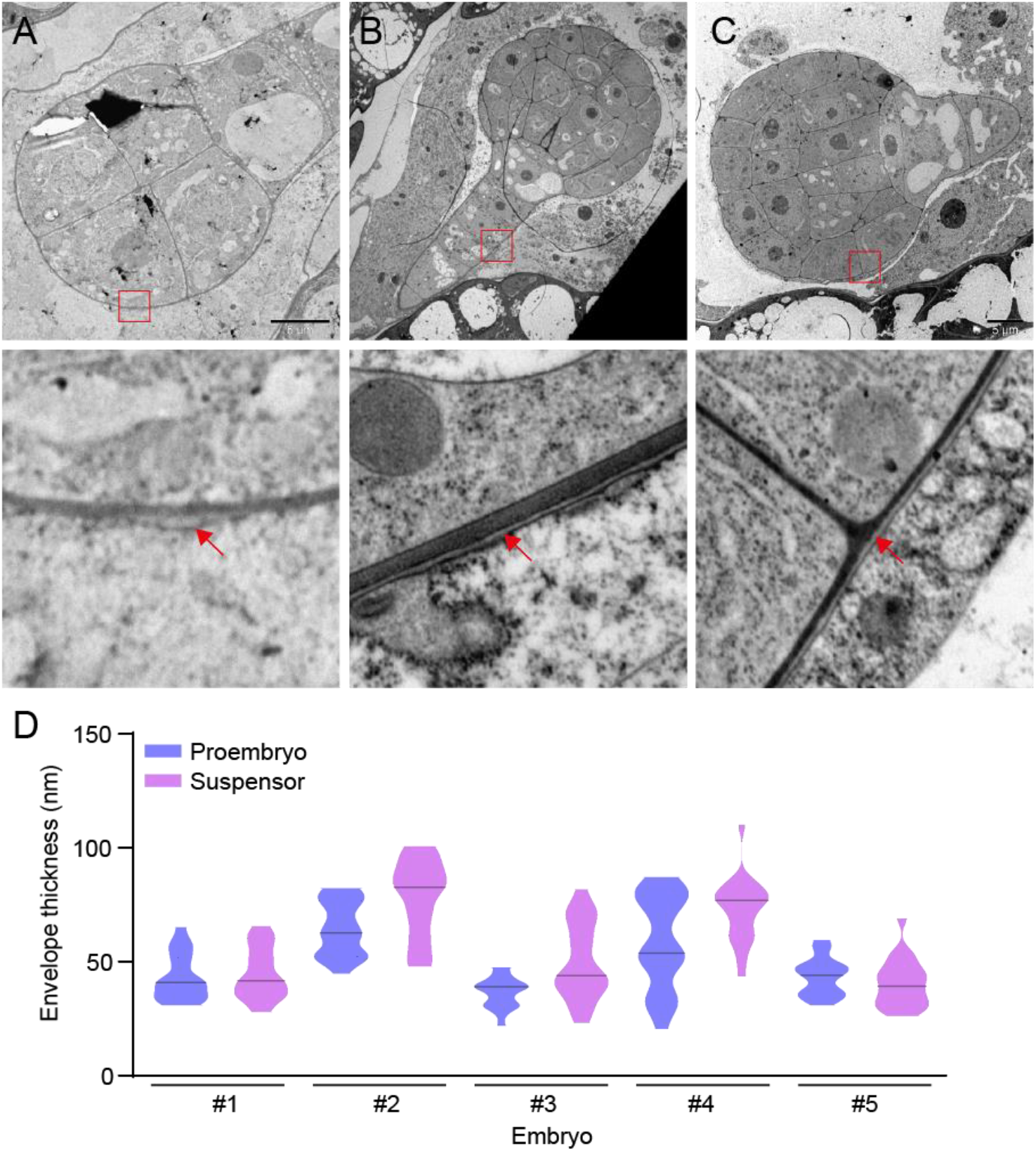
Ultrastructural analysis of the embryonic envelope. (A) TEM of Arabidopsis wild-type embryos at different stages, showing an electron-dense (white) layer in detail around the proembryo and the suspensor. Scale bar=XX μm. (B) Quantification of the thickness of the electron-dense (white) layer surround pro-embryo and endosperm, measured in sections along the proembryo and suspensor (n=5 embryos).

### Enzymatic liberation of the extra-embryonic envelope

From its appearance in TEM, it is difficult to tell what the envelope structure is made of. We first explored the possibility that this is a cell wall-like structure, and treated isolated embryos with a range of cell wall-degrading enzymes. Strikingly, none of the individually tested enzymes or mixtures (Cellulase, Pectinase, Hemicellulase, Driselase, Macerozyme), or combinations thereof led to visual dissociation of embryo cells (Supplementary Figure 1A), which would be an indication that any surface layer had degraded. When combining all these enzymes, however, we observed that a translucent layer parted from the embryo surface, creating a “halo” surrounding the embryo (Supplementary Figure 1A).

Following the same embryo over time showed that the structure was liberated after around 20 min of incubation (Figure 2A). Because the embryo shrinks as a consequence of prolonged enzyme treatment (Supplementary Figure 1B), the structure becomes more distinctly visible with incubation time, but does not visibly dissociate (Figure 2A). As seen for the features observed in TEM, this structure is found surrounding both the proembryo and the suspensor. When treating embryos isolated at different stages of development, we found the structure to exist from the earliest time point analyzed (8-cell stage) onward (Figure 2B). While we cannot formally demonstrate that the structure observed in TEM and upon enzyme treatment are the same, we interpret this as visualizations of the same extra-embryonic structure, that we term “embryonic envelope” (henceforth “envelope”). Since not even an aggressive mixture of enzymes that breaks a range of cell wall linkages can destroy the envelope, we conclude that its main structural component is not of cell wall polysaccharidic nature. The envelope does seem to be linked to the embryo cell walls, as enzymes that degrade the cell wall liberate it. Since only a complex mixture of enzymes is effective, the association of the envelope to the embryo cell walls is likely based on interactions or bonds with multiple cell wall polysaccharides.

**Figure 2.**
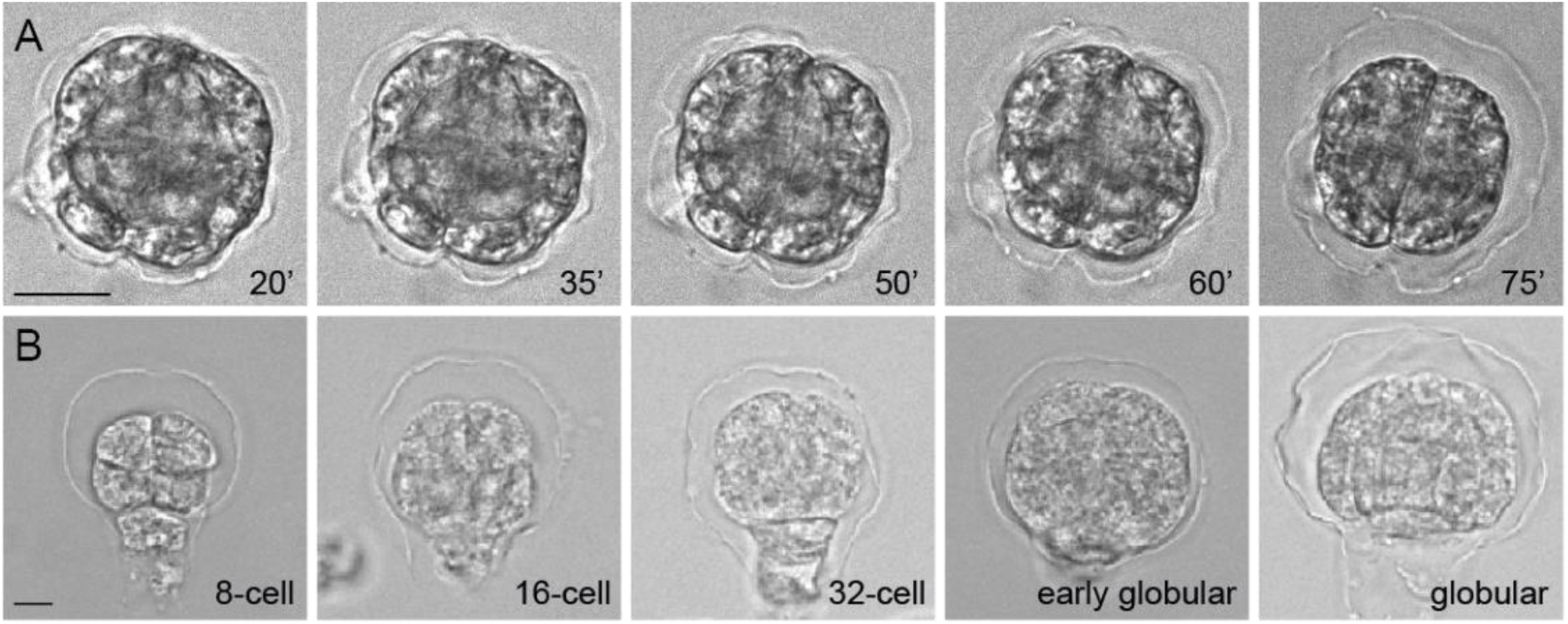
The embryonic envelope is present since early developmental stages. (A) Time course of the embryo envelope liberation (n=5 embryos). Scale bar=20 μm. (B) Different stages of *A. thaliana* embryos showing the envelope detached from its surface (n≥15 embryos per stage). Scale bar=10 μm.

### The envelope is distinct from the embryonic cuticle

It has previously been shown that cutinosomes carrying cuticular components start being deposited as patches at globular stage and the cuticle becomes a continuous structure at heart stage (Stępiński et al., 2017; Creff et al., 2019; Coen et al., 2019; Doll & Ingram, 2022). While the timing of cutinosome appearance and the early presence of the envelope suggests that the latter is not a cuticle-like structure, we tested this by analyzing mutants impaired in cuticle formation.

GASSHO1 and GASSHO2 receptor-like kinases have been shown to be required for embryonic cuticle deposition. *gso1/2* double mutants display a patchy cuticle and the embryos are physically attached to surrounding tissues (Tsuwamoto *et al*., 2008). We applied the same enzymatic treatment that liberates and allows visualizing the envelope to *gso1/2* embryos. We did not find a difference in the appearance of the envelope surrounding the proembryo and suspensor following enzyme treatment (Figure 3A, upper panels). Additionally, we tested a double mutant in two *Arabidopsis* acyltransferases, GPAT4 and GPAT8. *gpat4/8* double mutants are deficient in cutin biosynthesis and more susceptible to desiccation and pathogen infection (Li et al., 2007). Embryos from these mutant lines displayed the envelope after enzymatic treatment (Figure 3A, bottom panels). These results suggest that the embryo envelope is not an embryonic cuticle.

**Figure 3.**
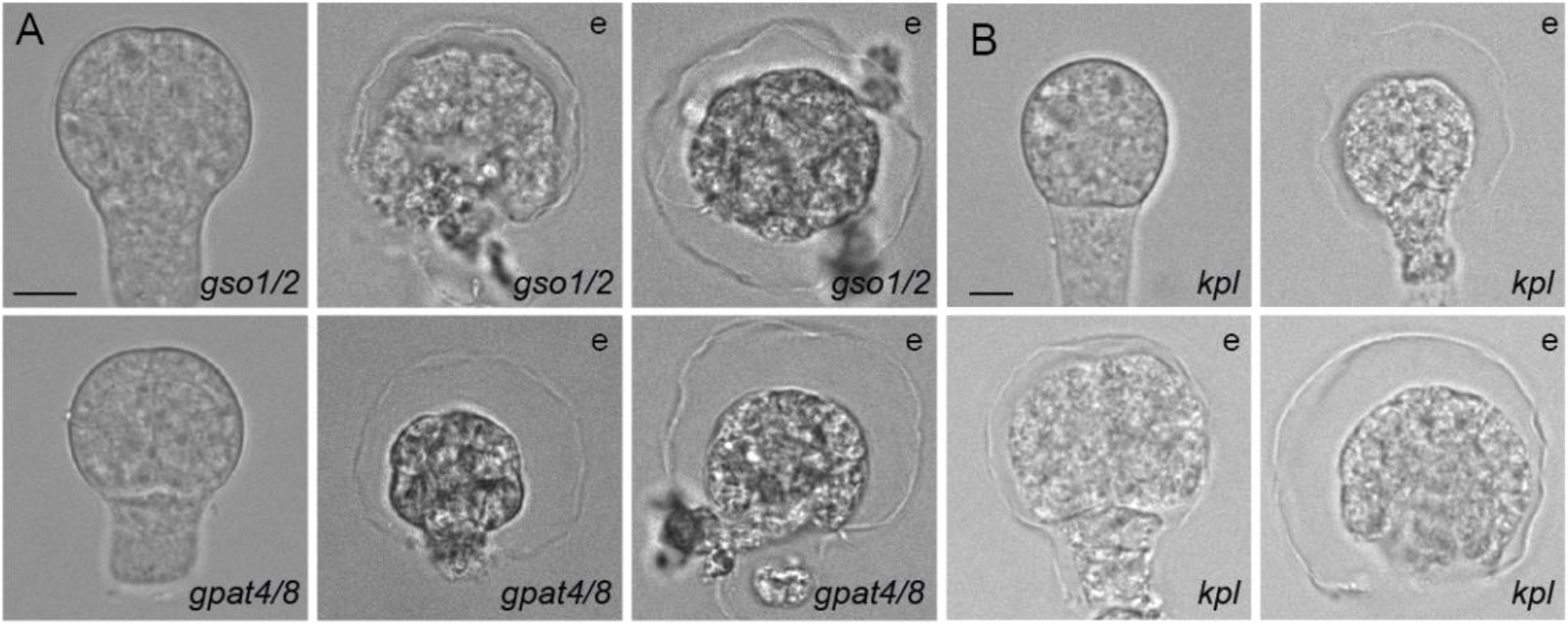
The envelope is not a cuticle and is an embryo-derived structure. (A) Embryos from cuticle-defective mutants, *gso1/2* and *gpat4/8*, showing the envelope (n≥40 embryos per genotype). (B) *kpl* embryos developed in absence of the endosperm, showing the envelope (n=40 embryos); e, after enzymatic treatment. Scale bar in all panels=10 μm. “e”, after enzymatic treatment.

### The envelope is embryo-derived

It has been shown that the material to build the embryonic cuticle is secreted by the embryo (Moussu et al., 2013) while the material to build the embryonic sheath is delivered by the endosperm (Moussu et al., 2017; Doll et al., 2020). We asked which tissue might generate the embryo envelope. To determine its origin, we tested the presence of the envelope in the *kpl* (*kokopelli*) mutants in which single fertilization events predominate (Ron *et al*., 2010). The phenotypes observed in *kpl* mutants include ovules that are unfertilized, ones that underwent double fertilization and contain both embryo and developing endosperm, as well as those with products of single fertilization, with developing embryo and unfertilized central cell or with only developing endosperm (Ron *et al*., 2010). These different classes can be distinguished by size. Based on seed size, we isolated only embryos developing in the absence of the endosperm and performed an enzymatic treatment. We observed detachment of the embryonic envelope in *kpl* mutants (Figure 3B), suggesting that the presence of endosperm is not required for envelope formation. While it is theoretically possible that the unfertilized central cell generates the envelope, we conclude that crosstalk between the endosperm and the embryo is not needed for the deposition of the envelope. These results indicate that the embryonic envelope is built from components secreted by the embryo. Importantly, this also defines the envelope as a unique structure, distinct from the endosperm-derived embryonic sheath (Moussu *et al*., 2017).

### The embryonic envelope is enriched in glycoproteins and lipids

The embryonic envelope is observed as a thin layer covering the embryo surface and it can be partially detached from it by enzymatic treatment. The integrity of the envelope upon treatment with cell wall-degrading enzymes suggests it is not a cell wall, and mutant analysis excludes that it is a cuticle. To investigate the composition of the envelope we tested a range of established fluorescent dyes and antibodies. Since the staining with such dyes and antibodies at the surface of intact embryos would not allow differentiating between the envelope and underlying structures, we stained embryos that had first been treated with wall-degrading enzymes to release the envelope.

First, we tested dyes that stain common components of the cell wall. Calcofluor white marks the primary cell wall polysaccharide cellulose (Wood 1980; Mori et al., 1996; Herburger et al., 2016) and, consistent with the results from enzymatic treatment, did not stain the embryonic envelope (Figure 4A). Aniline blue stains callose, a polysaccharide that is abundant in newly formed walls and plasmodesmata (Currier, 1957; Alexander, 1987; Zavaliev & Epal, 2015). No signal was observed in the envelope (Figure 4A). We also checked for the presence of pectins by using LM20 (binds to methyl esterified HG) and LM19 (binds to unesterified HG). No signal was detected on the surface of the embryos indicating that pectic polysaccharides are not present in this structure (Supplementary Figure 2). In contrast, the post-embryonic root surface was stained by both LM19 and LM20 antibodies (Supplementary Figure 3).

**Figure 4.**
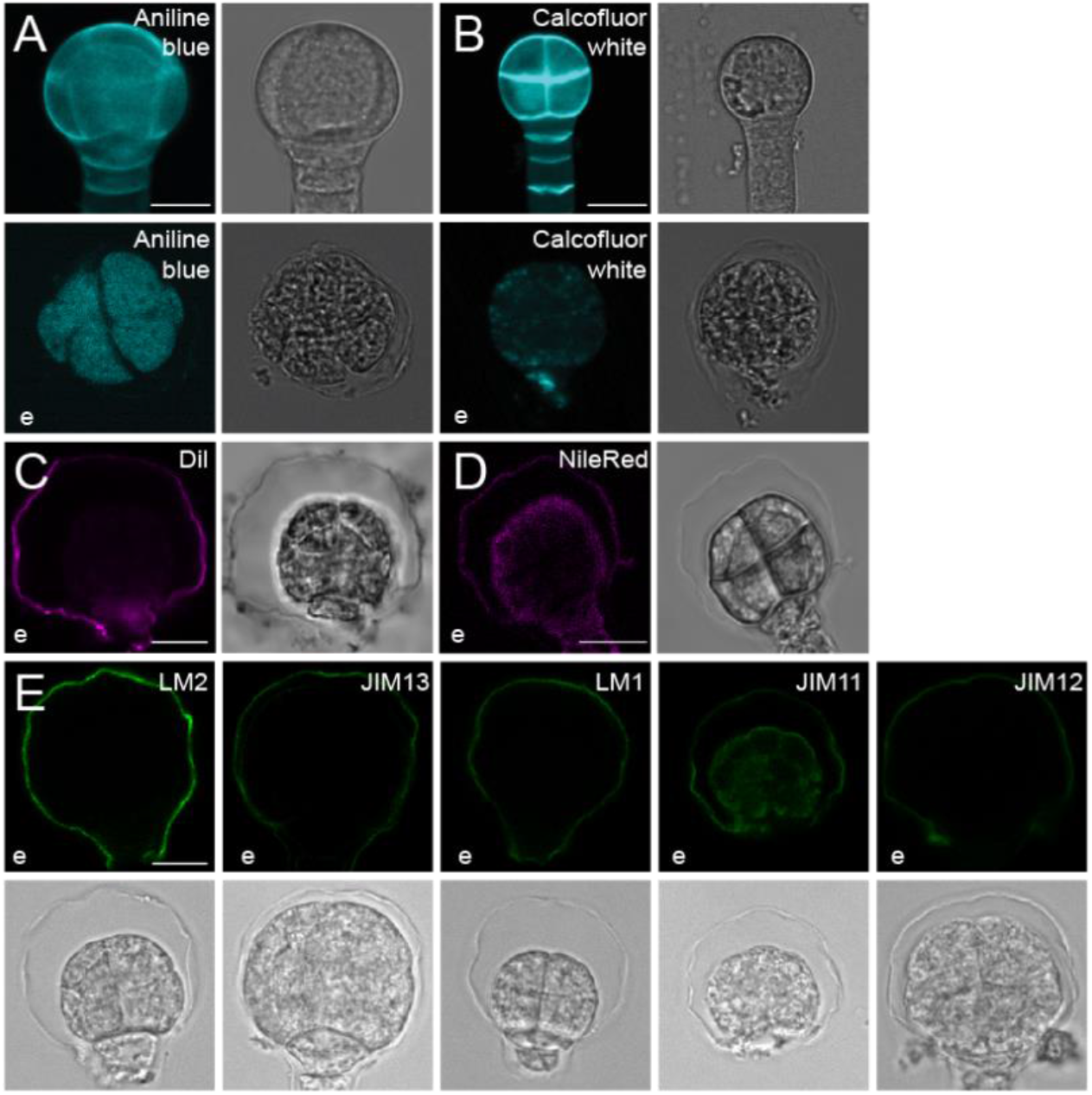
Embryonic envelope composition. (A-B) Calcofluor white (A) and aniline blue (B) staining of wild-type embryos (n≥50 embryos per staining). (C-D) Wild-type embryos stained with DIL (C) and NileRed (D) (n≥8 embryos per staining). (E) Immunolabeling of wild-type embryos (n≥30 embryos per staining) with LM2, JIM13, LM1, LM11 and LM12 antibodies. Scale bar=10 μm in all panels. “e”, after enzymatic treatment.

We next used the lipid dyes DIL and NileRed, that have been used before to stain different plant tissues such as roots and leaves and in protoplasts (Diaz et al., 2008; Blachutzik et al., 2012; Li et al., 2015). Both dyes generated a clear fluorescent signal in the embryonic envelope (Figure 4C-D), suggesting that lipids are a component of this structure.

In addition to polysaccharides and lipids, plant cells expose a rich landscape of extracellular proteins. Among these, two classes of *O*-glycoproteins are abundantly found in plant tissues: Extensins (EXTs) and Arabinogalactan Proteins (AGPs) (Kieliszewski et al., 1994; Kieliszewski et al., 2010; Ellis et al., 2010; Showalter et al., 2010). Importantly, the embryonic sheath is decorated by antibodies LM1 and JIM12 that label EXTs. We used a set of antibodies that label the glycosyl chains on hydroxylated prolines in EXTs. LM1, JIM11 and JIM12 each recognize a fully glycosylated chain on EXTs (Willats & Knox, 2003; Knox et al., 2008; Moller et al., 2008). We used a whole-mount immunostaining method aimed at labeling the surface layer. As this method does not involve permeabilization steps, we expect little to no staining of the embryo itself, except if the envelope is ruptured, and antibodies can penetrate the embryo. All three antibodies clearly stained the envelope. While LM1 and JIM12 appeared specific to the envelope, JIM11 also stained the embryo (Figure 4E). Again, given that the method was designed to address surface labeling, we do not interpret this differential staining of the embryo.

We next used two antibodies that detect branched glycosyl groups on hydroxylated prolines in AGPs. LM2 and JIM13 bind to beta-linked glucuronic acid and (beta)GlcA1→3(alpha)GalA1→2Rha epitopes, respectively (Willats & Knox, 2003; Knox et al., 2008; Moller et al., 2008). We found both antibodies to label the envelope (Figure 4E). Thus, the envelope clearly stains positively for both EXTs and AGPs.

To determine whether we could correlate envelope composition to structure, we labeled *gso1/2* and *gpat4/8* mutant embryos with the same panel of AGP and Extensin antibodies. While the envelope is present in these mutants (Figure 3A), there are defects in cuticle and sheath formation, and the latter in part labels for the same components (Doll et al., 2020). All antibodies still labelled the envelope in the *gso1/*2 double mutant (Supplementary Figure 3). In the *gpat4/8* double mutant, however, we found that the signal for each of the Extensin antibodies was undetectable (Supplementary Figure 3), despite clear presence of the envelope in transmitted light images. This suggests that the EXTs that are labeled by this panel of antibodies are not a critical component of the envelope. It also identifies AGPs as a likely key component of the envelope.

### The envelope has properties of a polymeric structure

A remarkable feature of both AGPs and EXTs is that these proteins can be chemically crosslinked (Kjellbom et al., 1997; Brady et al., 1997; Brady et al., 1998), forming polymeric structures. Given that we identified these proteins as likely components of the envelope, we asked if the envelope has properties that are consistent with crosslinked protein. To this end, we performed scanning electron microscopy (SEM) on isolated embryos that were either untreated or treated with cell wall-degrading enzymes. In control conditions (no enzymatic treatment) we observed the presence of a smooth continuous structure surrounding the entire embryo and suspensor. Clearly, dehydration of the embryo during processing created a deflated embryo and wrinkled surface (Figure 5A), which was decorated by a continuous structure following the entire wrinkled surface. We interpret this surface structure to be the envelope. As expected, the surface structure was comparable in *gso1/2* and *kpl* mutants (Figure 5B-C). To probe the properties of the envelope, we analyzed the enzyme-treated wild-type embryos. While the overall structure and morphology of enzyme-treated embryos was affected by the treatment, we found the surface structure to persist (Figure 5D). In places where embryo cells had come apart during the treatment or processing, however, we observed thread-like filaments that connected the separated cells (Figure 5D’-D’’). Detailed imaging of these areas revealed strings of globular structures that are 35 nm in diameter. The characteristic diameter for individual globular proteins is 5 nm (Lobanov et al., 2008), indicating that the features we observed are unlikely to be linear protein filaments.

**Figure 5.**
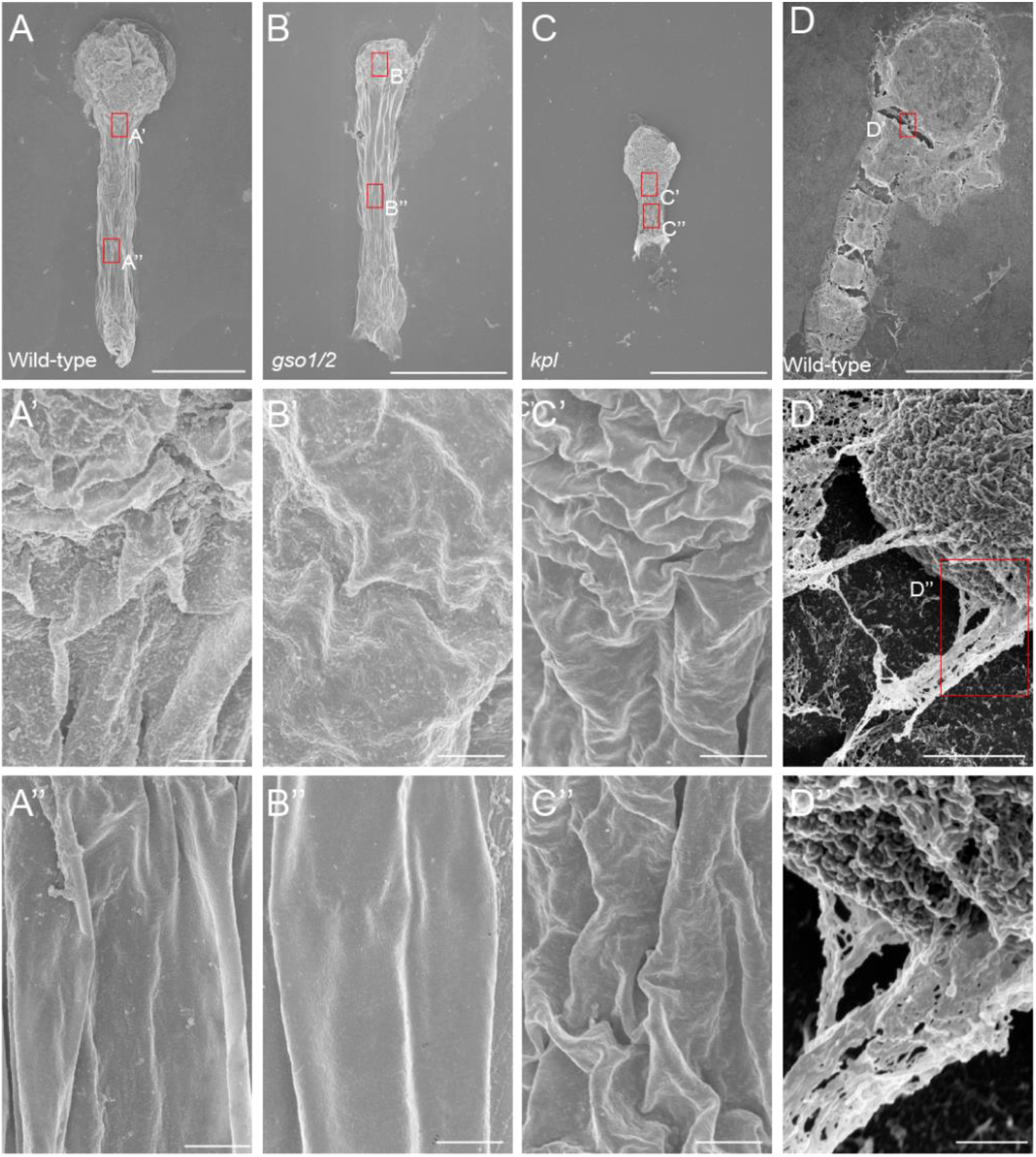
SEM on *A. thaliana* embryos showing the surface structure of the envelope. (A-C) SEM on wild-type (A), *gso1/2* (B) and *kpl* (C) embryos without enzymatic treatment. (A), (B) and (C) panels are overview images of entire embryos, (A’), (B’) and (C’) represent magnifications of surface of the proembryo, and (A’’), (B’’) and (C’’) are magnifications of suspensor surfaces. (D) SEM on Arabidopsis wild-type embryo after enzymatic treatment. (D’) and (D’’) represent magnifications of boxed areas in (D). Scale bars are 20 μm (A,B,C), 30 μm (D), 1 μm (A’, A’’, B’, B’’, C’, C’’) and 500 nm (D’, D’’) (n≥3 embryos for each genotype and treatment).

### Epidermal identity is required for embryonic envelope biogenesis

According to our results, the components needed to build the envelope are secreted by the embryo itself, but there are no known or suspected regulators of its production. However, given that the structure is formed on the surface, it is possible that its biogenesis is controlled by factors related to the epidermal identity of the surface-exposed cells. We therefore tested if defects in epidermal cell fate specification could affect the formation of the envelope. We performed enzymatic treatment on embryos from *pdf2 atml1* mutants. Since these double mutants are not viable or fertile as homozygotes, we used segregating plants that were homozygous for one and heterozygous for the other mutation (*pdf2 -/- atml1 +/-* and *pdf2 +/- atml1 -/-)*. We could observe the presence of the embryonic envelope in the mutants (Figure 6A). Interestingly however, the envelope appeared more damaged by the enzymatic treatment, showing discontinuities in its structure (Figure 6A-B) that were not observed in wild type embryos. This can be observed more in detail after the immunolabeling assays, in which a discontinuous fluorescent signal is observed (Figure 6B). In these immunolabeling assays, we found that the main protein components, AGPs and EXTs, are still present in mutant envelopes. To investigate if the surface of the envelope shows any evident defects we performed SEM on embryos from *pdf2 atml1* mutants but no clear differences were observed (Figure 6C). This could indicate that there are factors downstream the epidermal fate pathway affecting the composition of the envelope, more prone to break after the enzymatic treatment. Strikingly, we noticed that embryo cells appeared to dissociate from one another upon enzymatic treatment in the *pdf2 atml1* mutant (e.g Figure 6), a phenomenon that was never observed in wild-type embryos (e.g. Figure 2). Thus, the altered properties of the envelope likely render it more permeable to cell wall-degrading enzymes.

**Figure 6.**
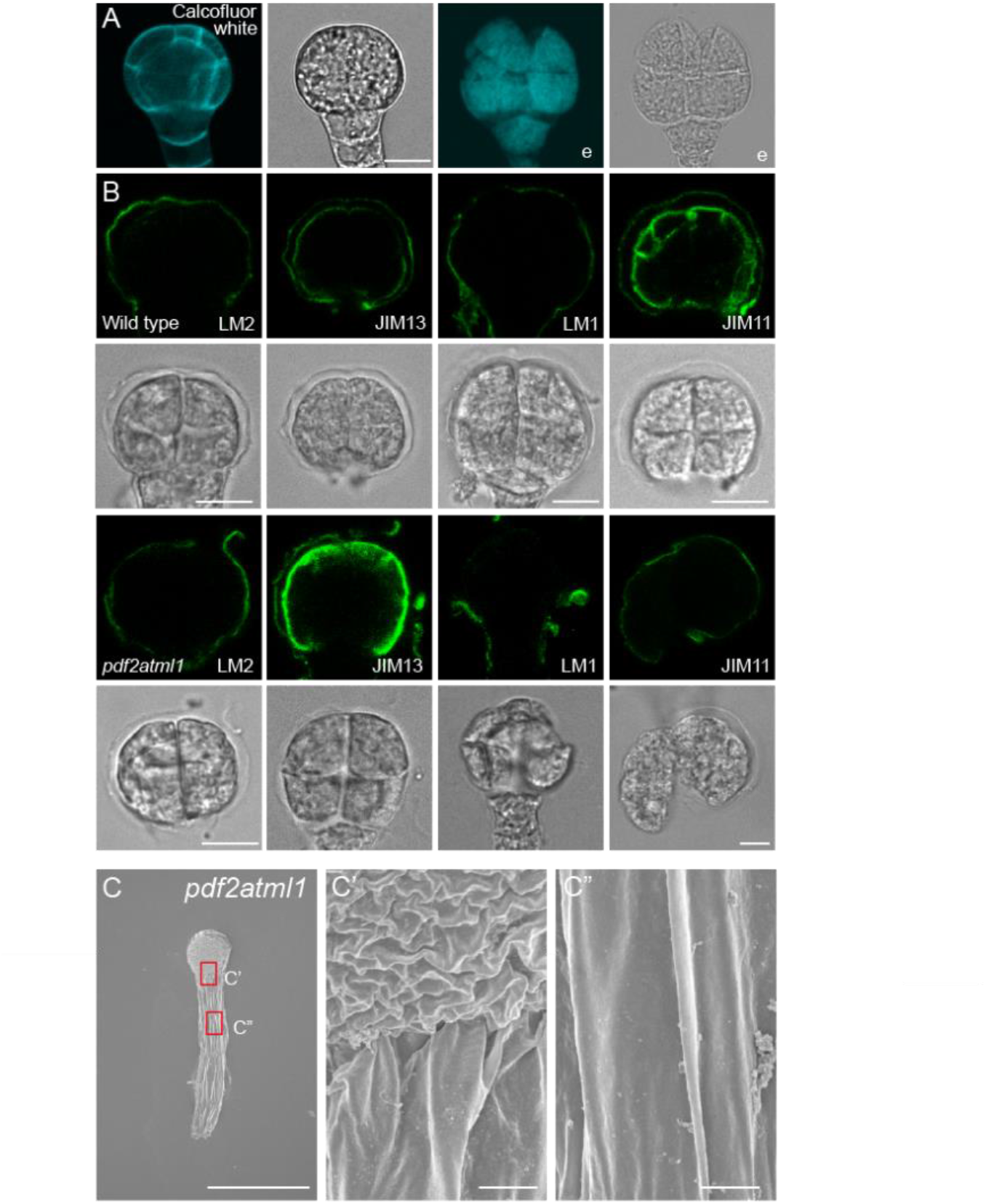
Defects in epidermal cell fate specification affect the formation of the envelope. (A) *pdf2 atml1* embryos stained with calcofluor white either without or with (“e”) enzymatic treatment (n=30 embryos). Scale bar=10 μm. (B) Immunolabeling of wild-type (top rows) and *pdf2 atml1* embryos with LM2, JIM13, LM1 and JIM11 antibodies after enzymatic treatment (n=50 embryos per genotype and antibody). Scale bar= 10 μm. (C) SEM of *pdf2 atml1* embryos in overview (C) and magnified details (C’ and C’’) (n=4 embryos). Scale bar= 30 μm (C) and 1 μm (C’ and C’’).

## Discussion

We have found a previously unreported thin membrane-like structure on the surface of early *A. thaliana* embryos. This embryonic envelope appears at early stages of development and it can be detected until late globular stage. To elucidate if this structure could be an early embryonic cuticle, we analyzed *gpat4/8* and *gso1/2* embryos since these mutants are defective in cutin synthesis and cuticle deposition, respectively (Tsuwamoto *et al*., 2008; Li et al., 2007). Embryos from both mutants displayed the thin continuous layer, showing that this structure is not the cuticle but the embryonic envelope. Another feature that distinguishes the envelope from the embryonic cuticle is that the former is found on the surface of the entire embryo while the latter is absent on the suspensor until early and late heart stage (Szczuka et al., 2003). Moreover, *kpl* embryos developing in the absence of an endosperm display the envelope, showing that a crosstalk between the embryo and the endosperm is not required for the production of this structure. Our results show that the envelope is composed of crosslinked *O*-glycoproteins and lipids. These biomolecules could give the embryo envelope its properties, being flexible enough to accommodate changes during embryo growth and development. Moreover, analysis of *pdf2 atml1* embryos shows that specification of epidermal cells is required for the proper production of the envelope since in those mutants this structure behaves differently after enzymatic treatment when compared to wild type. Although the components present remain largely the same, the embryonic envelope in these mutants seems to be more easily digested by the enzymes.

So far, we have not been able to assess the biological function of this structure, given the absence of a mutant that eliminates the envelope altogether. Raman imaging showed spectral peaks consistent with a proteinaceaous nature, but brief trypsin digestion of isolated embryos did not lead to the identification of major surface proteins (not shown). Therefore, the exact composition and key protein constituent of the envelope remains elusive, precluding genetic analysis. We can propose two hypotheses that will require deeper scrutiny in the future. First, the embryonic envelope could serve the same function as the sheath does in later developmental stages: forming an apoplastic diffusion barrier that both chemically and physically separates the developing embryo from the endosperm to allow both developmental programs to occur without unwanted crosstalk. Along the same lines, the envelope may serve as a template on which the cuticle and embryonic sheath are later assembled. Second, given the likely crosslinked proteinaceous nature of the envelope, the envelope could also serve a mechanical role. Plant developmental processes, in particular tissue patterning -which is crucial during embryogenesis (Palovaara et al., 2017; Harnvanichvech et al., 2021; Dresselhaus & Jurgens, 2021) - require a precise balance between outward directed osmotic forces, which are usually counterbalanced by tensile stresses in the stiff plant cell walls (Ali et al., 2023). In early stages of embryogenesis, cell walls need to accommodate substantial growth and thus need to be plastic and malleable (Malinowski& Filipecki, 2002). The endosperm itself undergoes rapid developmental changes at the same time, which are associated with large decreases in turgor pressure (Fourquin *et al*., 2016), and thus also cannot continuously provide a mechanical counterforce to the action of the increasing embryonic turgor. The embryonic envelope could play a role in resolving this mechanical conflict. Later, at the heart stage and beyond, the deposition of the embryonic sheath by the endosperm starts, separating the embryo from the environment and preventing the developing tissues to stick to the peripheral endosperm (Doll *et al*., 2020). At this stage, embryonic cell walls will be more capable of withstanding their own turgor, and the endosperm itself has reached a more turgescent state (Fourquin *et al*., 2016) to offer an additional mechanical counterforce. Further analysis needs to be carried out to unravel the composition, nature and function, and phylogenetic distribution of the embryonic envelope.

## Materials and methods

### Plant material

*Arabidopsis thaliana* ecotype Columbia-0 (Col-0), *gpat4/8, gso1/2, kpl-2* and *pdf2-1 atml1-3* seeds were surface-sterilized and grown on ½ Murashige & Skoog (MS) medium supplemented with 1 % sucrose (w/v) and 0.8% plant agar (w/v) and stratified at 4 °C for two days. Then, seeds were grown vertically on plates under long-day conditions (16 h light, 8 h dark) at 22 °C for two weeks. Seedlings were transferred to soil and further grown at a constant temperature of 22 °C under long-day conditions.

### Embryo isolation

Siliques were placed on double sided tape and sliced open in 1X Phosphate-Buffered Saline (1X PBS). Ovules from ∼ 20 siliques were collected in 2 ml Protein LoBind Tubes containing 20 μl of 1X PBS. Embryos were isolated with microcapillaries (VacuTipII, Eppendorf) and micromanipulator (Eppendorf) under an inverted microscope (Carl Zeiss Microscopy) as described by Raissig (2013), and were washed once in 30 μl of 1x PBS.

### Enzymatic treatment

Embryos were incubated in 20 mM MES (pH 5.7) containing 0.4 M mannitol, 20 mM KCl, 2% (w/v) cellulase R10 (Duchefa cat C8001), 0.5% (w/v) macerozyme R10 (Duchefa cat M 8002), 0.05% (w/v) Pectinase (Sigma cat P2401), 0.05% (w/v) Hemicellulase (Sigma cat H2125), and 0.05% (w/v) Driselase (Sigma cat D9515) for 20 minutes.

### Embryo envelope staining

After the enzyme treatment, embryos were incubated with 0.1% (w/v) Calcofluor White or Aniline Blue solutions for 5 minutes, and subsequently washed once with 1x PBS. For lipid staining, embryos were incubated with 0.1% (w/v) 1,1’-Dioctadecyl-3,3,3’,3’-Tetramethylindocarbocyanine Perchlorate (DiIC_18_(3)) or Nile Red, and subsequently washed once with 1X PBS. For immunolabeling, embryos were incubated in 20 μl of 3% (w/v) milk protein in 1X PBS (MP/PBS) for 15 minutes to block non-specific binding sites. Embryos were then incubated with primary monoclonal antibodies diluted 1:10 in MP/PBS for 15 minutes, and subsequently washed twice with 1X PBS. After washing steps, embryos were incubated with anti-mouse-IgG linked to Fluorescein isothiocyanate (Sigma cat.) diluted 1:100 in MP/PBS for 15 minutes.

### Confocal microscopy

Images were acquired in 8-bit format using a Nikon C2 Confocal laser scanning microscope with 63x NA = 1.20 oil-immersion objective with pinhole 1.0 Airy unit. Calcofluor White, Aniline Blue, DiIC_18_(3) and Nile Red were excited with 405 nm laser. Anti IgG-FITC was imaged in a SP8 Leica confocal microscope with 63x NA = 1.20 water immersion objective with pinhole 1.0 Airy unit, using 488 laser and HyD SMD2 detector.

### Transmitted Electron Microscopy imaging

Details of the optimized procedure will be presented elsewhere (Daghma *et al*., in preparation). Briefly, immature seeds were high pressure-frozen and freeze-substituted then infiltrated with Spurr’s resin and samples were transferred into BEEM capsules (Agar Scientific Ltd., Stansted, United Kingdom) and polymerized at 70 °C for 24 hours. Polymerized blocks were trimmed in pyramid shape while the face of the block was trimmed in trapeze shape. Semi thin (2 μm) and ultra-thin (70 nm) sections were cut with an ultra-microtome (Leica Ultracut, Leica Microsystems, Bensheim, Germany). The ultra-thin sections were post-contrasted in a LEICA EM STAIN (Leica Microsystems, Vienna, Austria) with uranyl acetate (Polyscience Inc. Eppelheim, Germany) for 30 minutes followed by incubation in Reynolds’ lead citrate (SERVA, Heidlberg, Germany) for 90 seconds and subjected to transmission electron microscopy (JEOL JEM2100, Peabody, USA) at 2.00 kV.

### Scanning Electron Microscopy imaging

Embryos were isolated as previously described, placed on poly-lysine coated glass slides, and allowed to settle at room temperature for 1 hour without drying. Once the embryos were attached to the glass they were gently rinsed with 1X PBS pH=7.4. A droplet of 2.5% glutaraldehyde was placed on the sample and fixation was carried out for 1 hour at room temperature. Samples were rinsed 3 times with 1X PBS. Post-fixation was carried out with 1% osmium tetroxide in phosphate buffer for 1 hour at room temperature. Samples were rinsed 3 times with distilled water and dehydrated in an ethanol series (30%, 50%, 70%, 80%, 90%, 96% and 2 x 100% ethanol, each for 10 minutes). Samples were affixed to flat aluminum stubs using double-sided adhesive carbon tape. A 12-nm layer of tungsten was subsequently applied using a Leica EM SCD 500 sputter-coater. Finally, images were captured using an FEI Magellan 400 field-emission SEM at an acceleration voltage of 2.0 kV.

## Acknowledgments

We would like to thank Marcel Giesbers and Jelmer Vroom from the Wageningen Electron Microscopy Centre at Wageningen University for their help in post-fixation with Osmium tetroxide and critical point drying of the samples for SEM.

Mutant seeds were kindly provided by dr. Taku Takahashi (Okayama University), dr. Gwyneth Ingram (ENS Lyon) and dr. Daisuke Maruyama (Yokohama City University).

## Author contributions

Y.H and C.B. carried out experiments, analyzed results, prepared figures. D.E.S.D and H.v.d.K. carried out experiments and analyzed results. J.S. and D.W. planned and directed the project. Y.H., C.B., J.S. and D.W. wrote the manuscript with input from all authors.

## Supplementary Figures

**Supplementary Figure 1.**
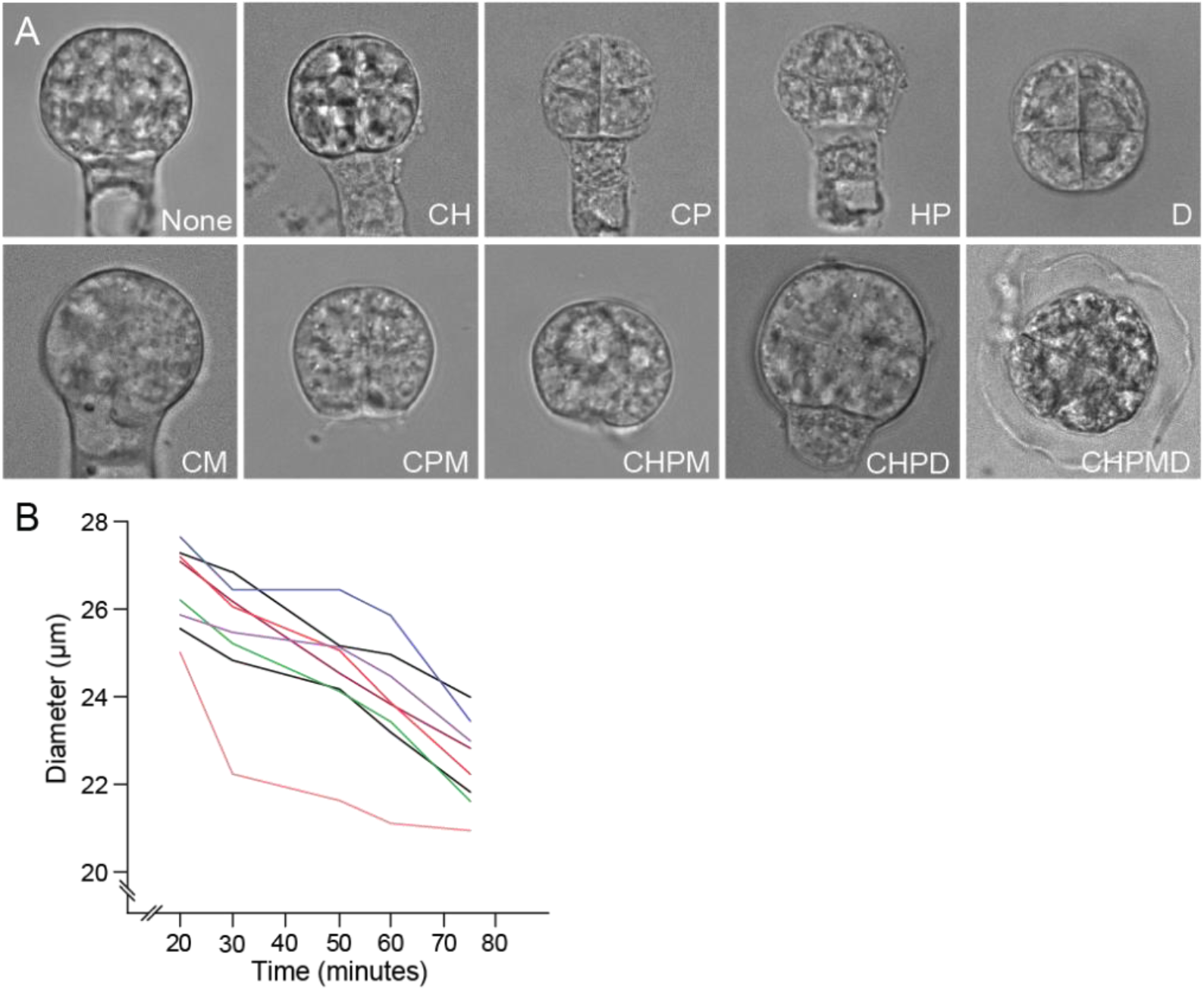
Wild type embryos subjected to different enzyme combinations. (A) Morphology of embryos treated with different enzymes or mixtures thereof for 20 minutes (C, cellulose; H, hemicellulase; P, pectinase; D, driselase; M, macerozyme). (B) Quantification of embryo diameter during incubation with CHPMD enzyme mixture. Each trace shows the diameter of a single embryo.

**Supplementary Figure 2.**
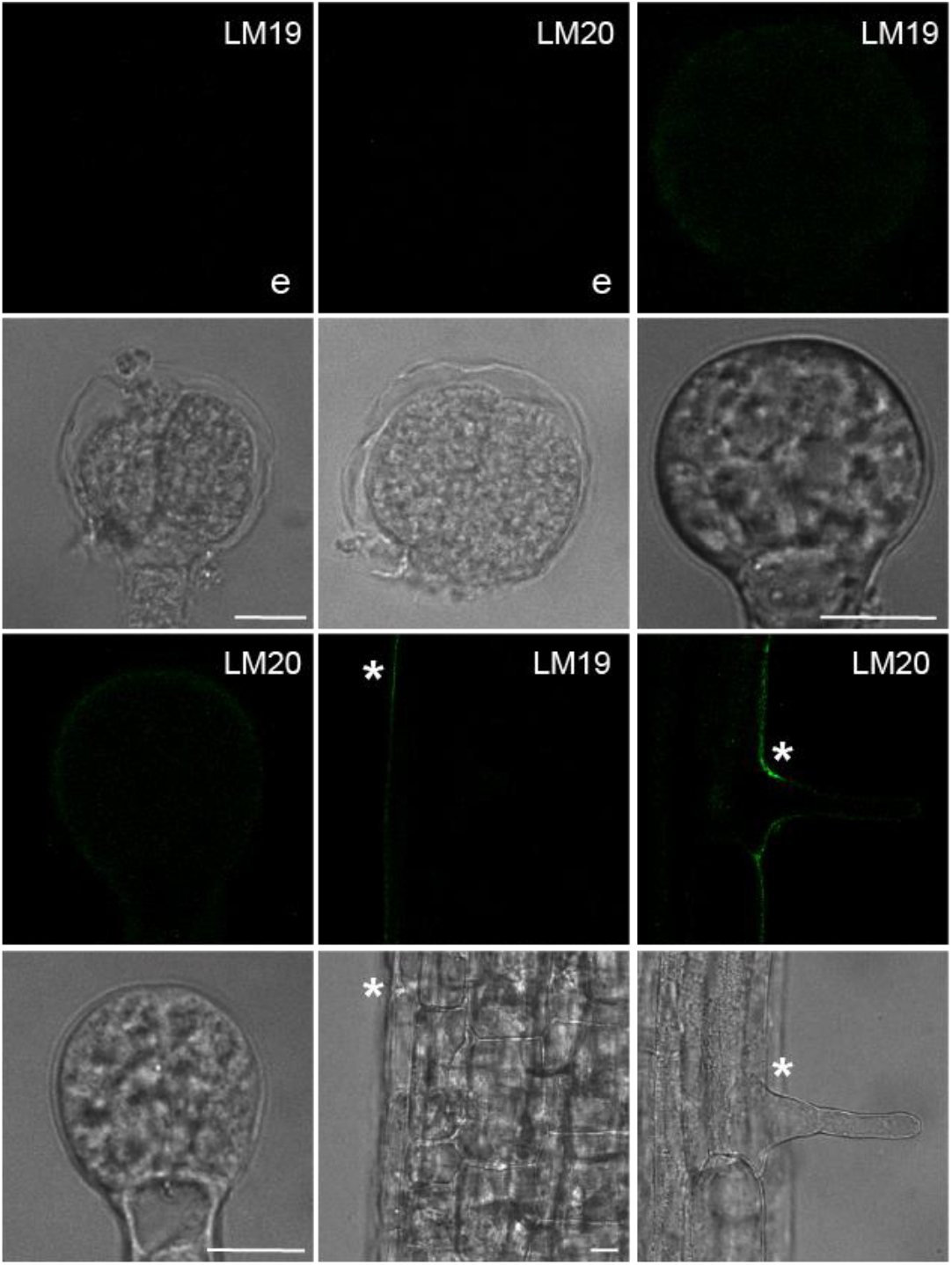
Pectin immunolabeling. Immunostaining of wild-type embryos (first 4 panels) and roots (last 2 panels) using LM19 and LM20 antibodies. “e”, after enzymatic treatment. Asterisks mark positive signal at root surface. Scale bar= 10 μm.

**Supplementary Figure 3.**
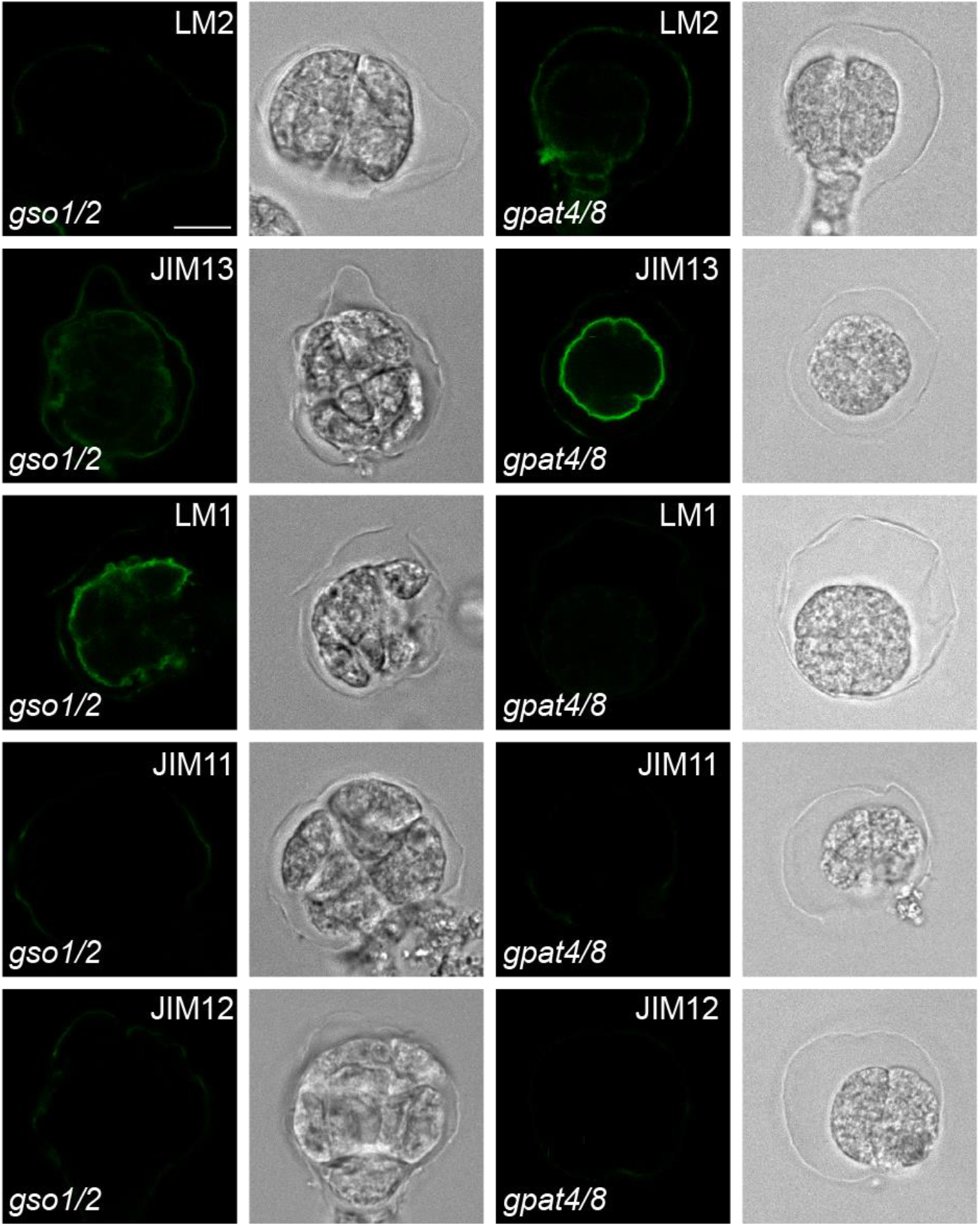
Mutant embryo envelope immunolabeling. Immunostaining of *gso1/2* and *gpat4/8* mutant embryos with LM2, JIM13, LM1, JIM11 and JIM12 antibodies after enzyme treatment. Scale bar= 10 μm.

